# Timing matters: age-dependent female receptivity and preference patterns

**DOI:** 10.64898/2026.06.23.734080

**Authors:** Lin Yan, Damian O. Elias

**Affiliations:** Department of Environmental Science, Policy and Management, University of California Berkeley, Berkeley, CA 94720, USA

**Keywords:** Female preference, Female receptivity, Jumping spiders, Reproductive timing, Mate choice

## Abstract

The timing of reproductive processes can influence how mating interactions shape reproductive strategies. We examined how female age correlates with receptivity and preference in the jumping spider *Habronattus formosus*, a system characterized by elaborate male courtship and strong female choice. To test how receptivity changes across the post-maturation period, we paired females of different ages with males and quantified male courtship displays and mating outcomes. Females were not receptive immediately after maturation and instead exhibited receptivity primarily 2–3 weeks after maturation. We further found that the full male courtship predicted mating success in older females, whereas only the second stage of the courtship predicted success in younger females. The delay in female receptivity is consistent with expectations for strong female choice systems, where females may have a greater opportunity to evaluate male courtship before mating. These results highlight the importance of age-dependent changes in female reproductive state and suggest that the timing of receptivity may shape how different components of male courtship influence mating success.

## Introduction

Sexual selection is a fundamental evolutionary process that shapes a significant portion of biodiversity (Andersson, 1994). Intra- (mate competition) and inter-sexual selection (mate choice) are the two primary mechanisms through which sexual selection typically operates and is thought to be dependent on (and in turn shape) an organism’s physiology, behavior, ecology, and evolution (Arnold, 1983; Hunt et al., 2009). One important yet often understudied aspect of sexual selection is the timing of reproductive behaviors like receptivity which is hypothesized to, among other things, influence the intensity and efficacy of intra- and intersexual selection. For example, when receptivity is predictable and/or synchronized, the strength and effectiveness of intrasexual competition is increased as animals are able to increase their fitness by restricting access to mates (Grafen & Ridley, 1983; Vellnow et al., 2020). This often leads to evolution of traits and behaviors associated with competition such as weapons and mate guarding (Daupagne et al., 2023; Hau et al., 2017). Alternatively, when receptivity is unpredictable and/or asynchronous the benefits of restricting access to mates is reduced often increasing the strength of intersexual selection thus leading to the evolution of traits used in mate assessment such as courtship signals.

A significant complication may also exist if the expression of reproductive traits and behaviors are not static and instead shift through time, likely influencing sexual selection dynamics (Carleial et al., 2024). Across animals, there is extensive evidence of age-related effects on reproductive behavior (Cotton et al., 2006; Kelly, 2018; Richard et al., 2005; Šmejkal et al., 2021). In reindeer, males’ reproductive effort is higher for older dominant males, likely because they reduce their effort in periods where females are not in oestrus, while young males maintain their effort across the breeding season even when females are not in oestrus (Tennenhouse et al., 2012). Age has been shown to impact choosiness and preference in females as well (Ah-King & Gowaty, 2016; Anjos-Duarte et al., 2011; Cotton et al., 2006; Dougherty, 2023; Ligout et al., 2012; Sarmiento-Ponce et al., 2021; Schneider et al., 2016; Tanner et al., 2019; Umbers et al., 2015; Wilgers & Hebets, 2012), the two major components of mate choice (G. G. Rosenthal, 2017). Sand goby females for example are increasingly choosy as the breeding season progresses (Forsgren, 1997). Age has also been seen to have the opposite effect on choosiness. For example, ovoviviparous cockroach females exhibit reduced choosiness and require shorter courtship durations as they age (Moore & Moore, 2001). Younger female crickets in habitats with high male density exhibit the most choosiness (Atwell & Wagner, 2014). Second, age has been shown to have effects on mate preferences. Coral damselfish maintained the same level of choosiness but reversed their preference functions when they got older (Wacker et al., 2016). Cichlid females shifted their preferences based on proximity to spawning (i.e. fertilization) and the presence of male hormones in the water from smaller to larger males (Kidd et al., 2013). Similarly, in crickets, younger females expressed strong preferences for attractive calls while older females did not (Gray, 1999). In another species of cricket (*Gryllus assimilis*), age explained 64% of the variation in the mate-finding behavior (i.e. phonotaxis) of females (Pacheco et al., 2013). While theory predicts that females should be more likely to accept mates as they age due to the increasing costs of mate searching and the urgency of reproduction (Cotter et al., 2011), studies suggest that shifts in mate choice may also stem from changes to sensory processing capabilities in choosers as they age (Kodric-Brown & Nicoletto, 2001; Ronald et al., 2012).

The Salticidae (jumping spiders) are a highly speciose family known for their diverse sexually selected traits and mating strategies. Like many arthropods, at the terminal molt, jumping spiders develop to adulthood as their adult reproductive anatomy: reproductive structures on pedipalps for males and epigyna for females, is expressed along with other secondary sexual characters. For many jumping spider species, males guard nests of soon to mature females (co-habitation) attempting to restrict access to other males (Jackson, 1980, 1986; Jackson & Hallas, 1986; Mendez et al., 2017). Males successfully defending terminal molt females gain fitness benefits as females are receptive immediately after maturation (Elias et al., 2014; McGinley et al., 2015). In other species, including species from the genus *Habronattus,* the subject of this study, co-habitation has not been observed (personal observation). *Habronattus* males are known for their complex multimodal displays that include color, motion, and vibratory songs (Bougie et al., 2024; Elias et al., 2003, 2012; Elias, Hebets, Hoy, et al., 2006; Maddison & Hedin, 2003; Maddison & McMahon, 2000; Maddison & Stratton, 1988; Rivera et al., 2021; Taylor et al., 2011). Female *Habronattus* tend to have relatively low mating rates (Blackburn & Maddison, 2015; Brandt et al., 2020; Elias et al., 2003, 2005; Elias, Hebets, & Hoy, 2006; Taylor & McGraw, 2013) and evidence suggests that courtship displays are essential to mating success (Elias et al., 2005; Elias, Hebets, & Hoy, 2006).

In this study, we examined the relationship between age, receptivity, and female courtship preferences in *Habronattus formosus*, a member of a species group with well-described complex courtship (*clypeatus* group; (Rivera et al., 2021)). Given the absence of mate guarding and co-habitation, we hypothesized that *H. formosus* females would not be receptive immediately after sexual maturation. Further, whether female preference pattern is consistent after sexual maturity is unknown in the group. We predicted that female preference patterns would shift with age. By exploring reproductive timing and its relationship to age in this species, we aimed to understand how sexual selection may be operating in this system and any change of female preference over time. We discussed what observed pattern associated with timing implies in the group.

## Materials and Methods

### Specimen Collection and Housing

Penultimate female and adult male *Habronattus formosus* were collected in Victorville, California in June 2022. Spiders were individually housed in 3x3x5 cm plastic containers (AMAC Plastic Products) with crinkled paper as environmental enrichment. One drop of water was provided per container each week along with 4-6 fruit flies or small crickets. Spiders were housed in the lab at 25 degrees Celsius with a 12:12 hour light cycle. Maturity checks on penultimate females were performed at least every 2 days to track maturity dates. We define age 0 as the day that a female spider is found with the presence of an epigynum for the first time, since the presence of epigynum anatomically enables the female to copulate with a male, regardless of behavior.

### Age assignment and mating trials

All mating trials were conducted between April and July 2022. Using a random number generator drawing from a uniform distribution in R, we assigned each female to an age (immature or 1 to 30 days post-maturation) at which the trial would be conducted. For each female, a randomly chosen sexually matured male was paired the day before the designated date for the trial. Spiders were fed the day before the trial. All males were only used once when possible (n=6/113 males used twice).

On the day of the trial, the female from each pair was allowed to acclimate in the experimental arena for a minimum of 5 minutes. The arena was made by stretching printer paper on a circular embroidery hoop (11cm diameter). A reflective piece of tape was placed in the middle of the arena to facilitate the recording of vibratory signals (Girard et al., 2011). An acetate ring (9cm diameter, 4.5 cm in height) was placed on top of the arena floor, with Vaseline applied on the interior of the ring to prevent the spiders from escaping. The arenas were illuminated by two LED light panels (160 LED CN-160, Neewar) and heated by a heat lamp (LF-13 Professional Series Dimmable Clamp Lamp, Zoo Med Laboratories) to ensure a controlled temperature of 31.04 ± 1.31 (mean ± sd) degree Celcius, as temperature can influence mating patterns (Brandt et al., 2018).

After 5 minutes of female acclimation, the male was dropped into the arena. Trials were run for 15 minutes, or until spiders copulated or were cannibalized (n=1). Trials were recorded with a mirrorless camera (Panasonic LUMIX GH4 4K Mirrorless Digital Camera) with a macro lens (Olympus M.Zuiko ED 60mm F/2.8 Macro). Vibratory signals were recorded using a laser vibrometer (Polytec PDV100). After the trial, both spiders were weighed using an Ohaus Analytical Plus scale. Mating trial videos were further analyzed using BORIS (Friard & Gamba, 2016). For each video, courtship duration, female aggressive display and copulation were marked. For courtship, we broke down the courtship display of males into two parts. During part 1, the introductory phase, the male sidles around the female in circles and gradually gets closer without any vibratory signals (Rivera et al., 2021). During part 2, the multimodal courtship phase, the male approaches the female in a straight line with various vibratory and visual displays. See detailed descriptions of the components in the multimodal phase of the courtship in (Rivera et al., 2021). Not all males that display part 1 courtship advance to part 2, providing an important source of variation in male responses to females.

### Statistical analysis

All statistical analyses were conducted in R version 4.1.3 (R Core Team 2022). We used a logistic regression to model female receptivity at different ages (days post maturity, age 0 is the date that females matured) with copulation success as a binary variable. Because sample sizes differed substantially among age groups, aggressive display duration was evaluated descriptively rather than statistically. Mean courtship durations of males across different age groups (immature, 0-10 days, 10-20 days and 20-30 days) regardless of mating success were compared with a linear model followed by emmeans and Tukey adjustments for pairwise posthoc comparisons. To evaluate how female preference patterns might change across age, we subset the dataset to exclude females that are younger than 10 days, since no copulation happened before that. We defined “young” (N = 35, age 10-20 days) and “old” (N = 32, age 20-30 days) females based on the threshold identified by the logistic regression model of female receptivity. We removed two outliers that had a mean introductory courtship duration of more than 150 seconds (more than 3 standard deviations above the mean). We again used logistic regression with copulation success as a binary response variable, and binary age group (young and old) and courtship measurements (mean duration of introductory or multimodal courtship, see above) as predictor variables. We selected the best model based on the lowest AIC scores, and significance levels of predictors were determined by type III ANOVA because of the presence of interaction terms.

## Results

### Female receptivity

Immature females never copulated with males. Nine (45%) of 20 males, however, did court immature females (9/20). In contrast, mature females mated during 32 (40%) of 80 trials. The earliest copulation occurred on day 10 after the molt to sexual maturation. The 50% likelihood for copulation occurred at 19.4 (95%CI = 16.3, 22.6) days post maturation (**Fig. 1a**).

**Figure 1.**
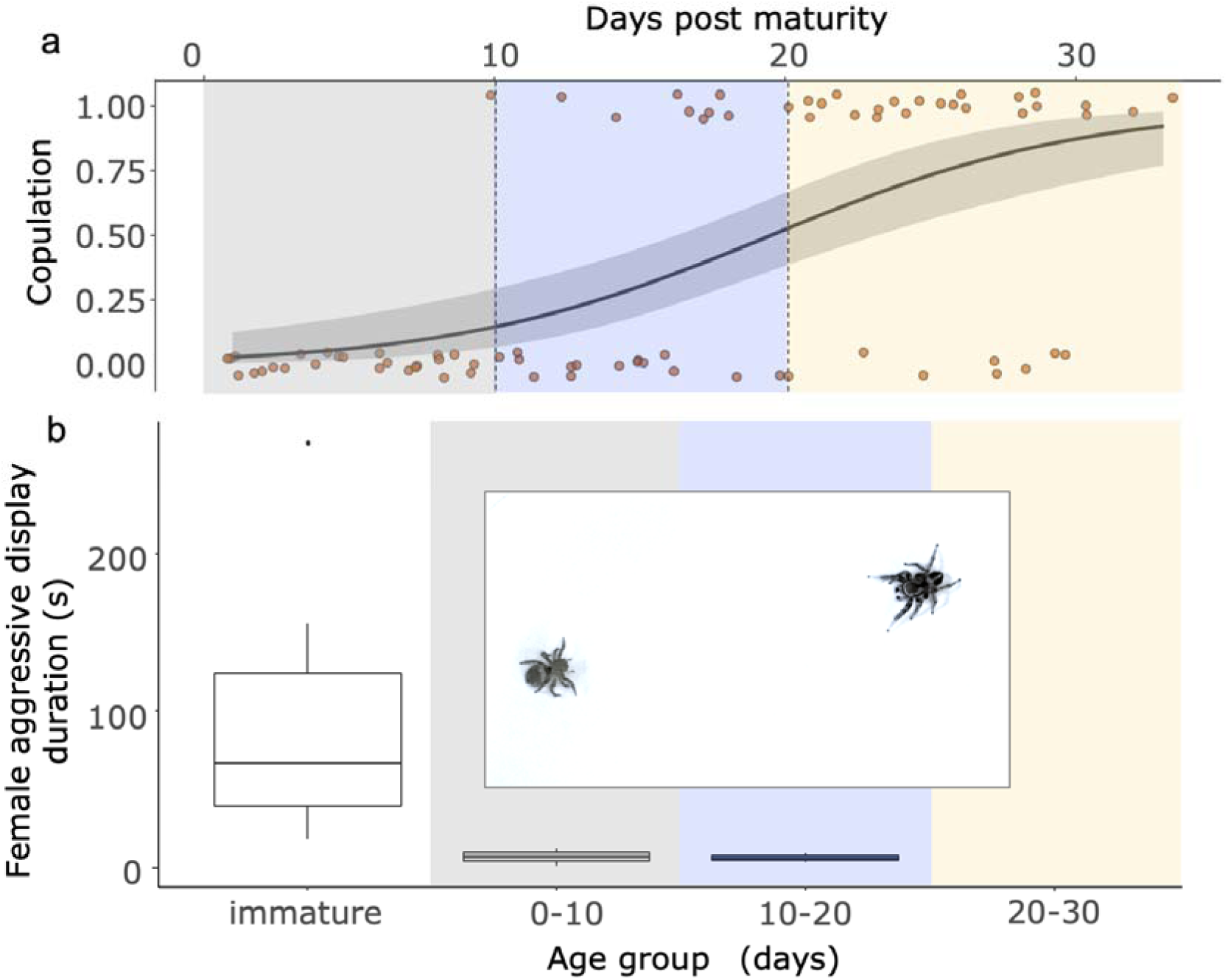
Female receptivity and aggressive display across age. **a:** Logistic regression model showing change of receptivity with age. Colored points are jittered along the y-axis for visualization, and shaded ribbons indicate 95% confidence intervals. **b:** Boxplots showing the total duration of female aggressive displays directed toward males across age groups. An example of this display is shown in the inset, with a subadult female displaying aggression on the left and an adult male attempting to court on the right. Age groups are indicated by the background shading in both panels.

### Female aggressive display

We discovered a female display that involves visually raising the first pair of legs outwards, with slight up and down movement, as well as a vibratory component of percussion (**Fig. 1b**). This behavior is predominantly found in immature females (17 out of 20) when interacting with males, but 3 individuals aged 0-10 days and 2 individuals aged 10-20 days were also found to perform the display briefly. No such behavior is observed in the 20-30 days age group (**Fig. 1b**).

### Male courtship across female age

In general, we found that male courtship duration is similar when courting females of age groups but different in others (**Fig. 2**). For the introductory courtship, mean courtship duration when courting immature females and 20-30 days old females are shorter than the mean duration when courting females of 0-10 days old (**Fig. 2a, Table S1**). For the multimodal courtship, no significant differences are observed in mean duration when courting females of adjacent age groups (**Fig. 2c, Table S2**). Notably, males exhibited similar courtship durations toward females in the 10–20 day and 20–30 day age groups during both phases.

**Figure 2.**
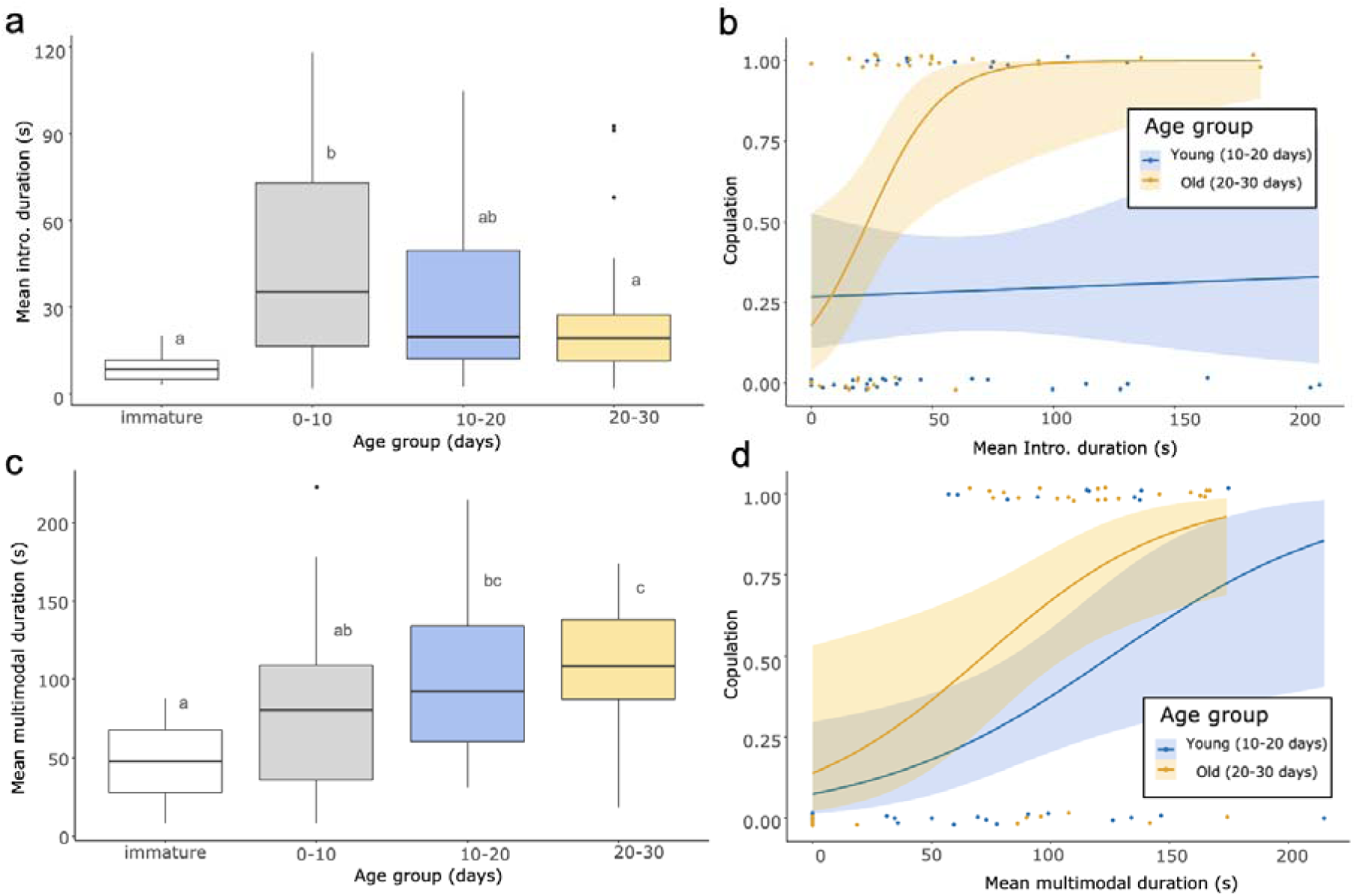
Male courtship duration and female preferences across female age groups. **a** and **c**: Boxplots of mean courtship duration of males when courting females at different age groups, letters indicate statistical significance. **b** and **d**: Logistic regression models of how male courtship display phases predict mating success. **a** and **b**: introductory display phase, **c** and **d**: multimodal display phase. Colored points are jittered along the y-axis for visualization, and shaded ribbons indicate 95% confidence intervals.

### Female preference patterns and age

We examined the relationship between introductory and multimodal courtship phases and mating across two age groups, young and old and found that preference patterns do change with age in *H. formosus*. The final model included age group, mean duration of introductory and mean duration of multimodal courtship phases, and the interaction of age and the mean duration of introductory courtship. The best model indicates that longer multimodal courtship plays a significant role in predicting copulation in both young and old females (Likelihood ratio chi-square (LR Chisq) = 15.1629, df = 1, p < 0.0001) (**Fig. 2d, Table S3**). However, the interaction between age and duration of introductory display also contributes significantly to the model that predicts copulation success (LR Chisq = 9.1631, df = 1, p = 0.0025). It further indicates that the introductory courtship predicts copulation differently in younger and older females: longer introductory courtship predicts copulation in older but not younger females (**Fig. 2b**).

## Discussion

In this study, we found that *Habronattus formosus* were not receptive immediately after maturation and instead, females became receptive two to three weeks after they matured to adulthood (**Fig. 1a**). We also found an aggressive display with visual and vibratory components predominantly in immature females. While aggressive behavior is common in female jumping spiders during courtship interactions (Jackson et al., 1998; Taylor et al., 2017; Vickers et al., 2025), this is the first case to our knowledge that stereotypical and multimodal display from females is described. Furthermore, we observed a shift in male courtship traits that predicted mating success as females aged. The entire complex male courtship (visual and multimodal phase) display predicted mating success in older females while only later parts of the courtship (multimodal phase) display predicted mating success in younger females.

### Evidence for delayed receptivity

As opposed to the predictable patterns of receptivity found in some jumping spider species where receptivity occurs immediately after maturation (Elias et al., 2014; Hoefler, 2007), we found that female receptivity was less predictable. No females were receptive in the first 10 days after maturation after which females became receptive over a period of about two weeks. Multiple factors may contribute to the variability in ages at which females first become receptive, such as variation in nutrition intake, genotypic differences, etc., and may not be adaptive. Regardless, the unpredictable gap between maturation and receptivity may provide females a way to prevent male manipulation/harrassment through an increase in costs and reduction of benefits to guarding females until they are receptive. This may provide opportunities for intersexual selection through female mate choice. In this scenario, males would invest in searching for receptive females and use elaborate courtship displays to access mating opportunities. The absence of mate guarding and male-male aggressive displays, in addition to the presence of complex multimodal courtship in *H. formosus*, are all consistent with this hypothesis. A delay in female receptivity after sexual maturation has been documented in a few other spider species in which female mate choice is important for reproductive interactions (Klein et al., 2012; Uetz & Norton, 2007), providing additional support for a connection between variable female maturation and elaborate male courtship displays.

### Delayed receptivity and complexity of courtship behavior

One potential consequence of this type of delay between maturation and receptivity is selection for greater complexity of male courtship displays. Delayed maturation may allow females the opportunity to sample courtship across multiple males, thereby allowing more informed choice of the males with which a female mates, since males still court females even when they are not receptive (Tuni & Berger-Tal, 2012). From the perspective of the females, the ability to both sample male densities and a large distribution of male courtship traits before they mate could potentially increase a female’s fitness if they were able to plastically respond to the availability of males as well as the male traits they experienced (Kappeler et al., 2023). It has been shown that redback black widows can respond to cues of male densities (Kasumovic et al., 2008; Kasumovic & Brooks, 2011; Lymbery et al., 2025; Scott et al., 2020).

### Comparisons with other species

Our finding that *H. formosus* females have a delayed receptivity aligns with anecdotal evidence in other spider systems in which female choice plays a dominant role. Some wolf spider females adjust their future mate choice patterns depending on the courtship they observed as juveniles (Hebets, 2003). These studies show that spiders can adjust their mate choice patterns depending on what they experience. Intriguingly, experimental studies on mate choice behavior in the wolf spider genera *Schizocosa* (e.g. (Hebets et al., 9/2013; Norton & Uetz, 2005; M. F. Rosenthal et al., 2018; Uetz & Norton, 2007)), the peacock jumping spiders (Maratus (Girard, 2017; Girard et al., 2018)), and other *Habronattus* (Brandt et al., 2020; Elias et al., 2005; Elias, Hebets, & Hoy, 2006), species/genera renowned for the complexity in their courtship displays, conduct experiments 2-3 weeks after maturation. While anecdotal, this does suggest a possibility that in spider species with complex courtship, there is a “critical period” where mating preferences develop. Future work examining this hypothesis, as well as what occurs during this period would help elucidate how mating preferences form. It further implies that delayed receptivity (and associated learning/experience-dependent mate choice) may be a feature of systems with diverse and complex communication during sexual selection events. In *Habronattus* species, preliminary observations suggest that females are predominantly monogamous. Delayed receptivity may reduce the costs associated with monogamy by allowing enough time to sample potential mates. Once again, similar findings in *Schizocosa (Norton & Uetz, 2005)* and *Maratus* (Girard et al., 2015) spiders suggest that this may be a feature of systems where females mate once and have complex communication. Future work should explore this hypothesis across different systems.

### Temporal variation in female preferences

When examining the relationship between male traits and mating success, we observed that in *H. formosus,* younger females selected males based only on a part of their display (multimodal courtship) while older females selected males based on the entire display (introductory courtship + multimodal courtship) (**Fig. 2b** and **2d**). The courtship display of *H. formosus* males follows a sequential pattern, progressing from an introductory display to multimodal displays showing a series of stereotyped motifs (Rivera et al., 2021). While successful matings involved all display stages, we found that the duration of the introductory displays did not influence copulation decisions in younger females, whereas both the introductory and multimodal display phases influenced mating success in older females. This suggests that older females assess the entirety of the display and thus incorporate more information into their decision-making processes than younger females do.

Comparisons of male courtship duration across female age groups indicated that males did not court receptive younger (10-20 days old) and older females (20-30 days old) differently (**Fig. 2a** and **2c**). As a result, the age-dependent differences in predictors of mating success are more likely attributable to changes in female preference than to changes in male courtship effort. Although male courtship duration varied among female age groups, this variation did not mirror the pattern of female receptivity. Female *H. formosus* begin to exhibit receptivity after 10 days post-maturation (**Fig. 1a**), however, neither the introductory nor multimodal courtship phase showed a corresponding shift before and after this threshold. This suggests that *H. formosus* males do not strongly adjust courtship duration based on female age alone. In contrast, male Schizocosa wolf spiders have been shown to respond differently to female silk cues associated with age and receptivity (Roberts & Uetz, 2005). Interestingly, despite the aggressive displays exhibited by most immature females during courtship interactions (**Fig. 1b**), the mean duration of the introductory display was not significantly different from that observed when courting females in the oldest age group (**Fig. 2a**). This pattern suggests that *H. formosus* males may invest similar courtship effort across females regardless of age or receptivity, or that selection for accurately assessing female receptivity is relatively weak. This interpretation is consistent with findings from other *Habronattus* species, in which males frequently court heterospecific females occurring in the same habitat (Taylor et al., 2017). Overall, courtship duration is the result of dynamic interactions between males and females, the observed variation likely reflects a combination of male and female behavioral responses. The mechanisms underlying these interactions remain to be disentangled.

While many studies have shown that experience affects mate choice (Dukas & Bailey, 2024; Hebets, 2003; Li et al., 2018; Stoffer & Uetz, 2023), the differences in female preference pattern observed in the present study are not due to prior mating experience, as all females were collected from the field as immatures and had never mated before the trials. This suggests the possibility that the shifts we observed were related to cognitive and/or physiological changes that occur with age (Kelly, 2018; Ronald et al., 2012; Rowell et al., 2021). Our findings may further suggest that younger females might not have the capacity to integrate information about the entire display like older females do. While some studies suggest that older females exhibit decreased sensory capacity in mating contexts (Ronald et al., 2012), this decline is typically observed after peak reproductive age. In contrast, our findings suggest that female preference patterns may continue to develop after sexual maturity, rather than simply declining with age. We suggest that this variation in female preferences could favor the evolution of more complex signaling like that found in *Habronattus* displays. While some theoretical models have suggested that over evolutionary time, courtship signals should evolve to be less complex (Iwasa & Pomiankowski, 1994), other authors have suggested that variation in female preferences may drive the evolution of more complex signals (Candolin, 2003; Jennions & Petrie, 1997; Reichert et al., 2017). We suggest that age-based variation in preferences may favor complex signals. Our results further highlight the importance of female age when measuring preference, as preferences may vary depending on their relationship to peak reproductive timing and shifts in the balance of selective forces.

Overall, our results suggest that reproductive timing is essential to understanding mating behavior. The timing between sexual maturity (morphological evidence) and sexual receptivity (behavioral evidence) may be associated with patterns of inter- and intra-sexual selection, and as a result, which traits or trait combinations are likely to evolve. We suggest that increases in the timing between sexual maturity and sexual receptivity are likely to favor the evolution of complex signals and/or preferences. Finally, we suggest that variation in the timing between sexual receptivity and sexual maturity across a population may favor the ability to plastically respond to relevant variation in potential reproductive partners. Regardless, understanding the timing of receptivity and how age impacts mate choice patterns is a critical, largely understudied aspect of mate choice. These types of studies are both needed to understand the dynamics of mate choice and to interpret behavioral data.

## Acknowledgments

We would like to thank the UC Berkeley Behavior group and the Elias lab for helpful discussions, especially Trinity Walls, Noah Leith, Jacob Gorneau and Sophie Hanson.We thank Annie Pho for assistance in specimen collection and undergraduate students in the Elias lab for animal care. This work was funded by a National Science Foundation grant to D.O.E. (DEB # 2327158).

---

**Supplementary Table S1.** Posthoc pairwise comparison of mean introductory courtship duration across female age groups

| pairs | estimate | SE | df | t.ratio | p.value |
| --- | --- | --- | --- | --- | --- |
| Immature - [0, 10) | -36.90 | 10.60 | 91 | -3.482 | **0.0042** |
| Immature - [10, 20) | -24.17 | 10.25 | 91 | -2.358 | 0.0928 |
| Immature - [20-30] | -17.08 | 10.44 | 91 | -1.635 | 0.3642 |
| [0, 10) - [10, 20) | 12.73 | 7.23 | 91 | 1.762 | 0.2986 |
| [0, 10) - [20, 30] | 19.82 | 7.50 | 91 | 2.642 | **0.0469** |
| [10, 20) - [20, 30] | 7.09 | 7.00 | 91 | 1.012 | 0.7428 |

**Supplementary Table S2.** Posthoc pairwise comparison of mean multimodal courtship duration across female age groups

| pairs | estimate | SE | df | t.ratio | p.value |
| --- | --- | --- | --- | --- | --- |
| [0, 10) - [10, 20) | -9.22 | 14.6 | 91 | -0.631 | 0.9219 |
| [0, 10) - [20, 30] | -42.61 | 15.2 | 91 | -2.809 | **0.0304** |
| [0-10) - Immature | 51.39 | 21.4 | 91 | 2.398 | 0.0847 |
| [10, 20) - [20, 30] | -33.39 | 14.2 | 91 | -2.358 | 0.0929 |
| [10-20) - Immature | 60.61 | 20.7 | 91 | 2.923 | **0.0223** |
| [20-30] - Immature | 94.00 | 21.1 | 91 | 4.450 | **0.0001** |

**Supplementary Table S3.** Model selection for logistic regression that predicts copulation. Intro_mean is the mean duration of introductory display, vib_mean is the mean duration of multimodal courtship, old is the age group (young/old) and copulation is a binary variable of whether copulation occurred.

| model | df | logLik | AICc | ΔAICc | weight |
| --- | --- | --- | --- | --- | --- |
| **copulation~vib_mean+old\*intro_mean** | 5 | -28.338 | 67.7 | 0.00 | 0.469 |
| cop~age_group*vib_mean*intro_mean | 8 | -24.873 | 68.2 | 0.57 | 0.353 |
| cop~age_group*vib_mean+age_group*intro_mean | 6 | -28.290 | 70.0 | 2.32 | 0.147 |
| cop~vib_mean | 2 | -35.123 | 74.4 | 6.77 | 0.016 |
| cop~vib_mean+intro_mean+age_group | 4 | -32.920 | 74.5 | 6.82 | 0.015 |
| cop~age_group | 2 | -42.109 | 88.4 | 20.75 | 0.000 |
| copulation~1 | 1 | -46.075 | 94.2 | 26.55 | 0.000 |

